# Neural processing of natural speech by adults with and without dyslexia: Evidence for atypical cortical decoding of speech information in the delta and theta EEG bands

**DOI:** 10.64898/2026.02.18.706607

**Authors:** Mahmoud Keshavarzi, Brian C. J. Moore, Usha Goswami

## Abstract

Neural oscillations in the delta (0.5–4 Hz) and theta (4–8 Hz) bands play a key role in tracking the temporal structure of speech. According to Temporal Sampling (TS) theory, dyslexia arises from atypical entrainment of these low-frequency oscillations to speech during infancy and childhood, which is particularly disruptive regarding phonological encoding. However, studies of adults with dyslexia have rarely examined both delta and theta cortical tracking under naturalistic listening conditions, and have not measured delta-band cortical tracking. Using EEG, here we focused on delta and theta band cortical tracking continuous natural speech by adults with and without dyslexia, applying a decoding analysis previously used with dyslexic children. Forty-eight English-speaking adults (24 dyslexic, 24 control) listened to a 16-minute continuous spoken narrative while EEG was recorded. Neural decoding of the speech envelope was quantified using backward multivariate Temporal Response Function (mTRF) models applied at two levels: a between-group analysis evaluating group-level differences in neural representation patterns, and a within-participant analysis assessing individual decoding accuracy. Cerebro-acoustic coherence was computed in parallel to provide a complementary measure of neural-speech synchronisation. Additional analyses examined band power, cross-frequency phase-amplitude coupling (PAC), and cross-frequency phase-phase coupling (PPC). Dyslexic adults exhibited less accurate delta- and theta-band decoding in the between-group analysis and reduced theta-band decoding accuracy in the within-participant analysis, alongside reduced coherence in both bands and increased delta-band power, particularly over the right temporal region. No group differences were found for PAC or PPC.

**Highlights:**

- Adults with dyslexia showed reduced delta- and theta-band speech decoding
- Cerebro-acoustic coherence was reduced in delta and theta bands in dyslexia group
- Delta-band power was increased in dyslexia, especially over right temporal region
- Cross-frequency coupling did not differ between adults with and without dyslexia

## 1. Introduction

Dyslexia is a developmental disorder characterised by difficulties in reading, writing, and spelling that cannot be attributed to low intelligence, inadequate educational opportunities, or obvious sensory impairments (Lyon et al., 2003). These difficulties with written language have been linked to impairments in phonological processing (Rack, 1994; Swan & Goswami, 1997; Snowling, 2000; Ramus, 2001). Oral phonological processing difficulties are characterised by a reduced ability to segment, identify, and manipulate sound elements within words. The Temporal Sampling (TS) theory (Goswami, 2011; 2015; 2022) provides a neurophysiological framework for understanding the phonological difficulties that characterise dyslexia across languages. TS theory proposes that dyslexia arises from atypical neural entrainment to slow temporal modulations in speech – particularly those at rates below 10 Hz. Low-frequency oscillations in the delta (0.5–4 Hz) and theta (4–8 Hz) bands are thought to support the tracking of syllabic and prosodic rhythms, allowing the brain to segment continuous speech into phonological units. Disruptions in these oscillatory mechanisms may lead to differently encoded phonological representations, thereby compromising the development of fluent reading and language skills.

For typical readers, neural oscillations in the delta and theta bands align closely with the temporal structure of speech, supporting efficient auditory and phonological processing. Delta-band neural oscillations track prosodic cues such as stress patterns and phrasal boundaries (Giraud & Poeppel, 2012; Gross et al., 2013). Theta-band oscillations track syllable onsets, potentially aiding the segmentation of speech into syllabic units (Di Liberto et al., 2015; Ding & Simon, 2014; Keshavarzi et al., 2020; Luo & Poeppel, 2007). This precise alignment between neural oscillations and speech rhythms is thought to underpin the parsing of spoken language and to play a fundamental role in developing phonological representations. For children with dyslexia, entrainment in these low-frequency bands (delta and theta) appears to be inconsistent or misaligned with the temporal structure of speech (Power et al., 2016; Molinaro et al., 2016; Mandke et al., 2022; Destoky et al., 2020, 2022, Keshavarzi et al., 2022 a; 2022b). Such misalignment may impair the extraction of prosodic and syllabic information, hindering speech parsing and affecting the automatic extraction of the linguistic hierarchy of stressed syllables, syllables, onset-rimes and phonemes.

A growing body of neuroimaging work has examined cortical tracking of natural speech using reconstruction-based models, such as backward Temporal Response Functions (TRFs; Crosse et al., 2016). Backward TRF models reconstruct features of the speech signal (e.g., the speech envelope) from neural responses, providing a quantitative measure of how the brain represents acoustic information. Reduced reconstruction (decoding) accuracy is interpreted as reflecting a less precise neural representation of speech. In dyslexia, such imprecision is thought to reflect atypical low-frequency entrainment, weakening the mapping between acoustic input and phonological representation of syllabic structure. Using electroencephalography (EEG), Keshavarzi et al. (2022a) reported reduced delta-band reconstruction of the speech amplitude envelope for dyslexic compared with control children. Using magnetoencephalography (MEG), Destoky et al. (2022) further demonstrated altered syllabic-rate (theta-band) cortical tracking in the right superior temporal gyrus of dyslexic children.

Another key neural measure relevant to dyslexia is cerebro-acoustic coherence, which is a measure of the synchronisation between neural activity and the temporal envelope of speech. Strong coherence reflects precise alignment of neural oscillations with acoustic cues – such as syllable stress patterns, word boundaries, and prosodic structure. Reduced coherence indicates weaker coupling between neural activity and speech rhythms, potentially disrupting the accurate development of phonological representations. Importantly, cerebro-acoustic coherence quantifies the strength of neural-speech synchronisation, but does not directly assess the fidelity with which acoustic information is represented in neural activity. Cerebro-acoustic coherence has been shown to be reduced in those with dyslexia, particularly for the delta band. Molinaro et al. (2016), for example, reported reduced delta-band coherence for both dyslexic children and dyslexic adults during sentence listening using relatively brief and segmented speech materials, with reduced synchronisation in the right auditory cortex and the left inferior frontal gyrus. While these findings demonstrate atypical low-frequency synchronisation, they leave open the question of whether dyslexic adults also show reduced precision in the neural representation of speech during sustained, naturalistic listening conditions. These disruptions in synchronisation may impair the extraction of the slow temporal cues that support the development of phonological awareness.

In contrast, cortical tracking studies of adults with dyslexia focused on the faster gamma band (30 Hz+) rather than on the delta band (Lehongre et al., 2011, 2013; Marchesotti et al., 2020). The processing of faster rates of temporal modulation is thought to be related to the extraction of phonemes from the speech stream, as exemplified initially by the ‘asymmetric sampling in time’ (AST) hypothesis (Poeppel, 2003). The AST hypothesis holds that the speech signal includes two timescales relevant to speech processing, a right-lateralised theta band timescale which relies on long temporal integration windows (150–250 ms) related to syllable-level information, and a left-lateralised gamma band temporal integration window (20–40 ms) related to phonetic information such as formant transitions. As individuals with dyslexia have difficulties with phoneme-level phonological processing as well as with prosodic and syllabic processing, Giraud and Poeppel (2012) proposed that they would exhibit atypical temporal sampling in the shorter temporal integration window of 20–40 ms. In an early study by Giraud’s group, Lehongre et al. (2011) investigated low gamma-band entrainment in dyslexic and neurotypical adult control participants with MEG, using a non-speech white noise stimulus amplitude modulated at rates ranging from 10 to 80 Hz. Dyslexic adults showed reduced low gamma-band entrainment in left auditory cortex relative to controls, and the strength of entrainment was correlated with measures of phonological processing and rapid naming. Further supporting the phonemic view, Lehongre et al. (2013) measured delta, theta and gamma oscillatory responses in adult dyslexics during passive viewing of an audiovisual movie that provided continuous speech input. The participants’ EEG was recorded at the same time as they underwent magnetic resonance imaging (MRI). Using a measure of correlation between participants’ oscillations and their MRI blood oxygen level dependent (BOLD) signal, intended to capture the degree to which different regions of interest oscillated in each frequency band, Lehongre et al. (2013) reported group differences in the lateralization of cortical responses for the gamma band only, with enhanced left-hemisphere correlations for control participants. Finally, to test whether gamma band activity was causally related to phoneme-level processing, Marchesotti et al. (2020) administered 20 minutes of transcranial alternating current stimulation (tACS) at 30 Hz to dyslexic adults to promote oscillatory responses in the gamma band. They reported improved performance following tACS in a subsequent phonological awareness task, although the effects disappeared an hour after stimulation. These findings suggest that by adulthood, atypical low gamma activity may be present alongside the delta and theta entrainment deficits highlighted by TS theory, impairing the processing of fine-grained phonetic cues that contribute to the development of phoneme awareness.

This developmental interpretation of the adult cortical tracking data is based on extensive behavioural research with children across languages, which has shown that phonemic awareness is a product of letter-sound learning and not a precursor of learning to read (Ziegler & Goswami, 2005). Accordingly, both TS theory and a phoneme-based theory may be correct, but may be applicable at different points in the developmental trajectory of reading acquisition. Indeed, the gamma-band differences between adults with and without dyslexia noted by Lehongre et al. (2011, 2013) may emerge as a consequence of the reduced reading experience that accompanies dyslexia. If repeated practice with letter-sound conversion facilitates phonemic awareness, then between the ages of 6–18 years children with dyslexia will accrue less practice than typically developing children which, over developmental time, could result in differences in gamma-band speech tracking.

The findings of Lehongre et al. (2011, 2013) were later replicated and extended by Lizarazu et al. (2021a), who used MEG to demonstrate that dyslexic adults exhibited significantly reduced neural phase-locking at 30 Hz (within the low-gamma range) to both speech and non-speech amplitude-modulated stimuli. The fact that reduced neural phase locking was not limited to speech suggests that by adulthood a domain-general impairment in auditory temporal sampling at phoneme-relevant rates may be present in individuals with dyslexia. Moreover, Lizarazu et al. (2021b) reported reduced neural responses to fast rises in amplitude in continuous speech for dyslexic adults. Although both dyslexic and non-dyslexic adults used such fast rises to phase-reset the neural oscillations tracking the speech signal, the dyslexic adults showed weaker phase locking values (PLVs) than the non-dyslexics in left auditory regions from ∼0.15 sec to ∼0.65 sec after such abrupt rises. Taken together, these results support the notion that disrupted phase synchronisation in the low-gamma band occurs for adults with dyslexia.

However, recent evidence supports the view that dyslexia in adults involves atypical neural entrainment across multiple temporal scales, not limited to the gamma band. Using a rhythmic audiovisual speech paradigm (repetition of the syllable “ba”), Keshavarzi et al. (2025) demonstrated that adults with dyslexia exhibited atypical neural entrainment in both low-frequency and higher-frequency ranges (see also Keshavarzi, 2025). While both dyslexic and control groups exhibited significant delta- and theta-band phase entrainment, the dyslexic group differed in preferred theta phase, indicating altered alignment to syllabic-rate information. Critically, whereas control participants showed robust beta- and low-gamma-band phase entrainment to rhythmic speech, entrainment at these temporal rates was absent for the dyslexic group.

Beyond neural decoding and coherence, atypical oscillatory power has also been reported for those with dyslexia. Band-power differences reflect the strength of activity within specific frequency ranges. They have been observed during both speech-listening and resting-state paradigms. Dyslexic individuals often show increased power in low-frequency (delta, theta) and beta bands. The functional significance of these power differences is not yet understood. They may reflect compensatory recruitment of additional resources during auditory processing tasks. Resting-state EEG studies of dyslexic children have shown increased delta and theta power in frontal and right temporal regions (Arns et al., 2007) and elevated theta power in frontal and temporal language regions (Papagiannopoulou & Lagopoulos, 2016). Using functional near-infrared spectroscopy (fNIRS) to measure blood flow, Cutini et al. (2016) observed increased right temporal responses to nonspeech stimuli amplitude modulated at a low rate (2 Hz). Consistent with these observations, Penolazzi et al. (2008) reported increased delta-band EEG activity in children with developmental dyslexia during linguistic processing tasks. Elevated delta power was associated with poorer linguistic performance, leading the authors to suggest that enhanced delta activity may serve as a neural marker of dysfunctional language processing. Task-based studies have also shown atypical beta-band power in individuals with dyslexia. Power et al. (2016) employed an EEG paradigm in which dyslexic and control children had to report specific words in semantically unpredictable sentences and observed significantly higher beta-band power for the dyslexic group. Similarly, Keshavarzi et al. (2024) used an audiovisual rhythmic paradigm involving the repetitive presentation of the syllable “ba” at 2 Hz and found significantly elevated beta-band power for dyslexic compared with control children.

In the current study, we investigated neural decoding for adults with and without dyslexia during natural speech listening by collecting EEG data. As adult EEG data are typically less noisy than child data, we were able to utilise some powerful analysis methods. Using backward mTRF models (Crosse et al., 2016), we first quantified low-frequency speech envelope decoding in the delta and theta bands. Two decoding approaches were employed: within-subject decoding and between-group decoding (Keshavarzi et al., 2022a). For within-subject decoding, a backward mTRF model was applied to each participant, and decoding accuracy was quantified by averaging performance scores across participants within each group. For between-group decoding, we trained normative backward models on a subset of the control group’s data and tested their generalisation on the remaining control participants as well as the dyslexic group. This approach enabled a direct comparison of decoding performance between dyslexic and control groups. Secondly, we calculated cerebro-acoustic coherence in the delta and theta frequency bands and band power in the delta, theta, beta, and low-gamma bands. Third, we computed cross frequency phase-amplitude coupling (PPC) and phase-phase coupling (PAC) measures. The PPC and PAC measures were computed across frequency pairs including delta-theta, delta-beta, theta-beta, delta-low gamma, and theta-low gamma, and separately for each group.

Based on TS theory and prior studies conducted using children with dyslexia, we anticipated that individuals with dyslexia would exhibit lower decoding accuracy than controls for the delta band, using both between-group and within-participant analyses. Consistent with TS theory and Keshavarzi et al. (2025), we also hypothesised that dyslexic participants would show less accurate theta-band decoding than controls. This prediction was also based on the prior observation that theta-band entrainment was positively correlated with reading efficiency in EEG studies with typically developing children (Power et al., 2012). On the basis of TS theory, we further hypothesised that cerebro-acoustic coherence for both the delta and theta bands would be lower for individuals with dyslexia than for control participants. Based on prior child dyslexia data, for the band-power analyses we predicted significant differences in delta-band power localised to the right temporal region (Arns et al., 2007; Penolazzi et al., 2008; Cutini et al., 2016) and in beta-band power across the whole brain (Keshavarzi et al., 2024) between the dyslexic and control groups. However, based on prior child data, we did not expect differences between the two groups for delta-beta PAC (Keshavarzi et al., 2024; Power et al., 2016). We had no specific hypotheses regarding PAC measures for delta-theta, delta-low gamma, theta-beta, and theta-low gamma interactions or for PPC measures involving delta-theta, delta-beta, delta-low gamma, theta-beta, and theta-low gamma oscillations across the groups.

## 2. Methods and Material

### 2.1. Participants

Forty-eight adult native English speakers took part in the study, comprising 24 individuals with dyslexia (16 female; mean age = 21.4 ± 2.5 (standard deviation) years; range = 18.3–29.1 years) and 24 control participants without dyslexia (16 female; mean age = 21.9 ± 2.6 years; range = 18.7–29.2 years). Participants with dyslexia were recruited through the Accessibility & Disability Resource Centre at the University of Cambridge, and all had a formal clinical diagnosis of dyslexia. Inclusion criteria required a confirmed diagnosis of dyslexia and the absence of additional learning difficulties such as autism spectrum disorder, dyspraxia, attention deficit hyperactivity disorder, or developmental language disorder. The absence of comorbid conditions was verified through participant self-report. In addition, all dyslexic participants scored at least one standard deviation below the control group mean on at least one subtest of the Test of Word Reading Efficiency – Second Edition (TOWRE-2; Torgesen et al., 2012). Control participants were matched to the dyslexic group as closely as possible for age and nonverbal IQ, measured using the Matrix Reasoning subtest (Wechsler, 2008; see Table 1). As shown in Table 1, there were no significant group differences in age or Matrix Reasoning scores. In contrast, the dyslexic group showed significantly poorer performance on standardised measures of reading fluency and phonemic decoding, assessed using the Sight Word Efficiency (TOWRE-SWE) and Phonemic Decoding Efficiency (TOWRE-PDE) subtests of the TOWRE-2, as well as slower performance on two versions of the Rapid Automatized Naming task (RAN 1 and RAN 2; Thomson et al., 2013; Richardson et al., 2004). Mathematical ability, assessed using the Test of Written Arithmetic (TAN; Wilkinson & Robertson, 2006), did not differ significantly between groups. All participants underwent pure-tone audiometry (250–8000 Hz) for both ears and were required to have hearing thresholds ≤ 20 dB HL at all frequencies. The same cohort of participants was tested by Keshavarzi et al. (2025). Written informed consent was obtained from all participants in accordance with the Declaration of Helsinki. Ethical review was undertaken by the Psychology Research Ethics Committee at the University of Cambridge, which gave a favourable opinion for the study.

**Table 1.** Mean scores (standard deviations) for age and behavioural measures for the control and dyslexic groups. Measures were age, single word reading accuracy and nonword decoding (TOWRE-SWE, TOWRE-PDE), rapid automatized naming (RAN 1 and RAN 2), nonverbal IQ (Matrix Reasoning), and mathematical ability (TAN). Group differences were assessed using two-tailed Wilcoxon rank-sum tests.

|  | Control | Dyslexia | Wilcoxon rank-sum test outcome |
| --- | --- | --- | --- |
| Age | 21.4 (2.5) | 21.9 (2.6) | $p = 0.55$ |
| TOWRE-SWE | 118 (12.9) | 94 (9.7) | $p = 1.1 \times 10^{-6}$ |
| TOWRE-PDE | 119 (8.9) | 98 (8.3) | $p = 5.8 \times 10^{-8}$ |
| RAN 1 (ms) | 21 (2.0) | 25 (4.1) | $p = 2.9 \times 10^{-4}$ |
| RAN 2 (ms) | 27 (2.7) | 33 (7.0) | $p = 7.9 \times 10^{-5}$ |
| Matrix Reasoning | 15 (1.6) | 15 (1.8) | $p = 0.53$ |
| TAN | 76 (10) | 72 (11.6) | $p = 0.22$ |

### 2.2. Experimental set up and stimuli

Participants were seated in an electrically shielded, sound-proof room to minimise external noise and electromagnetic interference. Auditory stimuli were delivered binaurally via insert earphones (ER-2, Etymotic Research) at a sampling rate of 44.1 kHz, while EEG was recorded simultaneously with a 1-kHz sampling rate using a 128-channel HydroCel Geodesic Sensor Net (Electrical Geodesics Inc.). Each participant listened to a 16-minute spoken excerpt from *The Iron Man: A Children’s Story in Five Nights* by Ted Hughes, presented as continuous, natural speech to approximate real-world listening conditions. This paradigm was designed to examine sustained neural entrainment and auditory processing during naturalistic speech comprehension. The same speech material (the first 10 minutes) had been used previously with children by Keshavarzi et al. (2022a). During listening, participants were instructed to remain still and fixate on a red cross (“+”) displayed at the centre of a monitor to minimise eye movement artefacts. EEG data were continuously recorded throughout the session to capture the brain’s real-time response to the unfolding narrative.

### 2.3. EEG data pre-processing and analyses

EEG data were referenced to the Cz electrode and band-pass filtered between 0.2 and 42 Hz using a zero-phase finite impulse response (FIR) filter. The data were then downsampled to 200 Hz. Noisy channels were identified manually and interpolated using MNE-Python (Gramfort et al., 2013). Independent component analysis (ICA) was performed to remove artefacts, components being identified based on scalp topographies, power spectra, and time-domain waveforms. For decoding-related analyses, additional filtering was implemented in MATLAB. Each frequency band was extracted using a combination of third-order high-pass and low-pass Butterworth filters designed with the *fdesign()* function. Specifically, the cut-offs were set at 0.5 Hz and 4 Hz for the delta band and at 4 Hz and 8 Hz for the theta band. Filtering was applied to the EEG channels using the *filtfilthd()* function. For cerebro-acoustic coherence analyses the cut-off frequencies were set to 0.5 and 8 Hz. For PAC and PPC analyses, the delta, theta, beta, and low-gamma bands were extracted using MNE-Python. Power analyses were performed on unfiltered data. After band extraction, all datasets were further downsampled to 100 Hz to reduce computational load.

### 2.4. Stimulus-reconstruction using backward TRF model

Neural decoding of speech was quantified using a backward mTRF model (Crosse et al., 2016). This linear model reconstructs a stimulus feature (here speech envelope) from neural activity, estimating how well cortical responses track the temporal dynamics of speech. The reconstructed envelope, 𝑠(𝑡), was computed as:

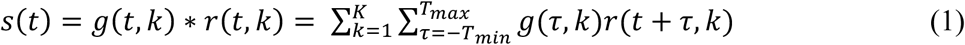

where 𝑔(𝜏, 𝑘) represents the decoder weight linking EEG channel 𝑘 to the stimulus envelope for time lag 𝜏, and 𝑟(𝑡, 𝑘) is the neural response at channel 𝑘 and time *t*. Lags from 𝑇_()*_ = 0 ms to 𝑇_(+,_= 400 ms were used, capturing the temporal window of cortical tracking.

To prevent overfitting and enhance model generalisability, the backward TRF model incorporated Tikhonov regularisation (Tikhonov, 1977), which penalises large model coefficients to balance model complexity and fit. The ridge parameter (λ) controls the strength of this penalty: smaller values permit greater model flexibility, whereas larger values enforce stronger regularisation to avoid overfitting. Model validation was performed using a leave-one-out cross-validation procedure (implemented via the *mTRFcrossval()* function from the TRF Toolbox). In this approach, data from each trial were sequentially excluded from training and used for testing, while data fort the remaining trials were used to fit the model. This procedure was repeated across all trials in the training set, so that each trial (in the training set) contributed to model validation. To identify the optimal regularisation parameter, eight candidate λ values (*λ* = 0.1, 1, …,10^6^) were evaluated. The value yielding the highest average Pearson correlation between the reconstructed and actual stimulus feature during cross-validation was selected as the optimal regularization parameter. This optimal value was subsequently used to train the final TRF model. Model performance was quantified as the Pearson correlation between the reconstructed and true envelopes, higher correlation values indicating better reconstruction of the envelope.

### 2.5. Within-participant backward TRF analysis

To assess the decoding accuracy of low-frequency components of the speech envelope (0.5–8 Hz) at the individual level, a separate backward TRF model was constructed for each participant. The speech envelope was obtained by computing the absolute value of the analytic signal derived from the Hilbert transform (*hilbert()* function, MATLAB) and subsequently band-pass filtering it between 0.5 and 8 Hz. EEG data were band-pass filtered separately in the delta and theta bands. This within-participant approach enabled evaluation of individual neural decoding accuracy and comparison between dyslexic and control participants. For each participant, both EEG data and the corresponding speech signal were segmented into 16 epochs of approximately one minute. Thirteen segments were used to train the TRF model, and the remaining three were reserved for testing, ensuring that model performance was evaluated on unseen data. After training, the participant-specific model was applied to the test data to reconstruct the speech envelope from EEG responses. Decoding accuracy was quantified as the Pearson correlation between the reconstructed and actual envelopes, computed separately for each neural band (delta and theta). Individual correlation values were averaged within each group to obtain group-level decoding scores, enabling direct comparison of neural speech decoding between adults with and without dyslexia. Figure 1 shows a schematic diagram of the analytical framework used for the within-participant backward TRF analysis.

**Figure 1.**
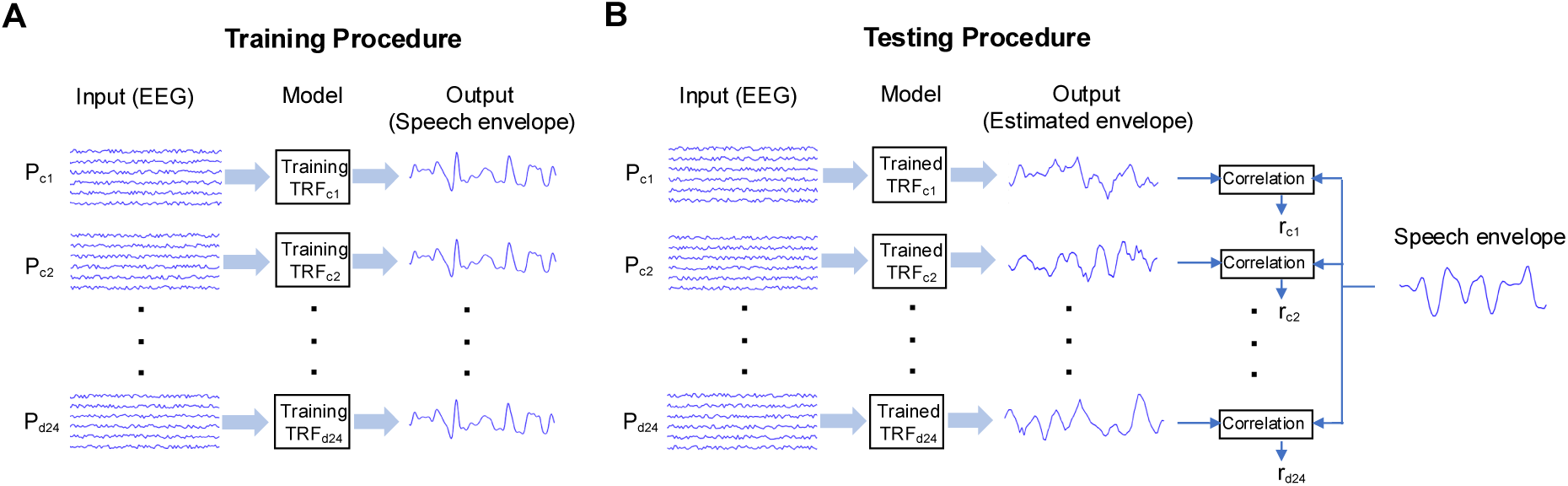
Schematic diagram of the framework used for the within-participant analysis. Panel A illustrates the procedure used to train the model and panel B shows the procedure used to test the model. *P_ci_* (*i* = 1, 2, …, 24): the *i*th control participant; *P_di_* (*i* = 1, 2, …, 24): the *i*th dyslexic participant; *r_ci_* (*i* = 1, 2, …, 24): decoding accuracy for the *i*th control participant; *r_di_* (*i* = 1, 2, …, 24): decoding accuracy for the *i*th dyslexic participant. This figure is reproduced with permission from Keshavarzi et al. (2022a).

### 2.6. Between-group backward TRF analysis

To evaluate group differences in the pattern of the neural representation and to simulate normative encoding patterns, a between-group backward TRF analysis was performed. EEG data from the control participants were used to construct a normative decoding model for each frequency band of interest (delta and theta). This approach provided a reference framework for assessing how closely the neural responses of individuals with dyslexia aligned with typical neural representations of speech. For each frequency band, a randomly selected subset of 12 control participants (Cₜᵣₐᵢₙ = 12) was used to train individual backward TRF models. These models were averaged to generate a single normative model representing typical decoding performance. The averaged model was then tested independently on the remaining 12 control participants (Cₜₑₛₜ = 12) and on all 24 dyslexic participants. Decoding accuracy for each test participant was quantified as the Pearson correlation between the reconstructed and actual speech envelopes. To ensure robustness, this procedure was repeated 100 times with random reassignment of participants to the training and test sets, generating 100 normative models. Mean decoding accuracies across iterations were computed separately for the control and dyslexic groups. This iterative approach minimised bias from any specific train-test split and enabled a stable estimate of group differences in the decoding of low-frequency speech information. Reduced decoding accuracy in the dyslexic group was interpreted as reflecting altered or less typical neural representations of the speech envelope. Figure 2 shows a schematic diagram of the framework used for the between-group backward TRF analysis.

**Figure 2.**
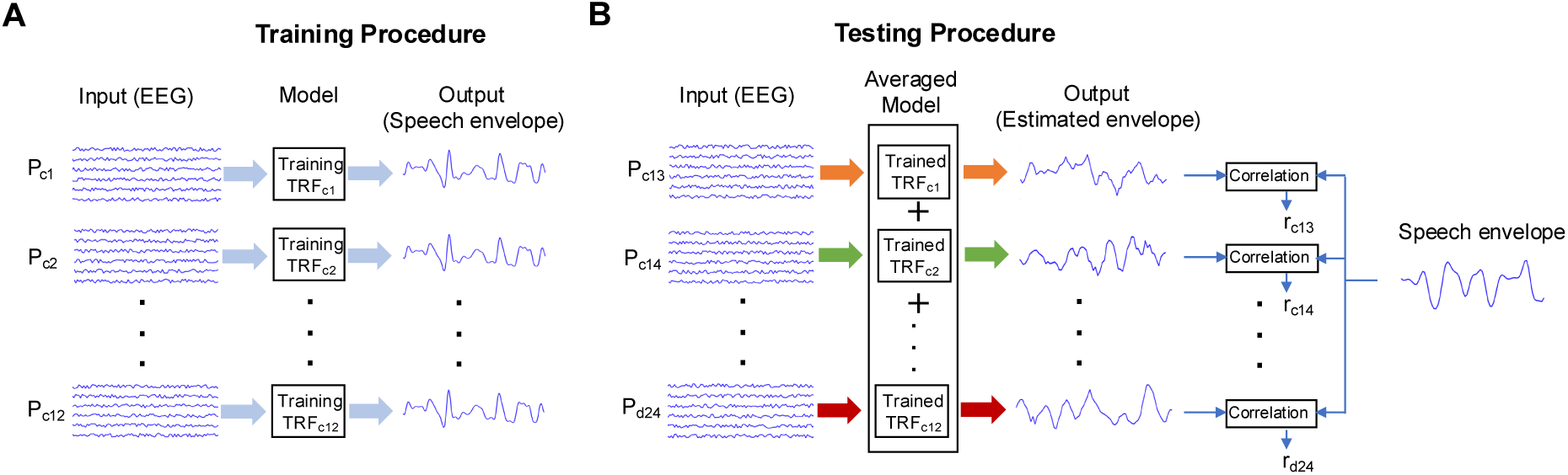
Schematic diagram of the model used for the between-group analysis. Panel A illustrates the procedure used to train the model and panel B illustrates the procedure used to test the model. *P_ci_* (*i* = 1, 2, …, 24): the *i*th control participant; *P_di_* (*i* = 1, 2, …, 24): the *i*th dyslexic participant; *r_ci_* (*i* = 13, 14, …, 24): decoding accuracy for the *i*th control participant; *r_di_* (*i* = 1, 2, …, 24): decoding accuracy for the *i*th dyslexic participant. This figure is reproduced with permission from Keshavarzi et al. (2022a).

### 2.7. Computation of chance level for backward TRF models

To assess the statistical significance of stimulus reconstruction accuracy obtained from both the within-participant and between-group backward TRF models, null models were generated for each frequency band of interest (delta and theta). These null models established data-driven chance-level decoding performance. For each iteration, EEG data from 12 randomly selected control participants were permuted across different story segments in the training set, thereby disrupting the temporal correspondence between EEG activity and the original speech envelope. Importantly, the testing segments were not permuted and remained aligned with the true speech signal. This ensured that model training was based on non-informative mappings, while evaluation was performed using the actual stimulus. Each null TRF model was trained on 13 permuted one-minute segments and tested on three unpermuted segments. The Pearson correlation between the reconstructed and actual envelopes was computed for each participant and frequency band, yielding a single null-model correlation per participant. The mean correlation across the 12 participants represented the group-level chance accuracy for that iteration. This process was repeated 100 times for each frequency band, with new random subsets of control participants in each iteration. The resulting distributions were used to construct probability density functions (PDFs), which defined the empirical chance level for evaluating statistical significance in the observed decoding results.

### 2.8. Computation of cerebro-acoustic coherence

To assess the coherence between neural activity and the speech envelope, cerebro-acoustic coherence was computed for each participant using EEG data (filtered between 0.5 and 8 Hz) and temporally aligned with the speech envelope (filtered between 0.5 and 8 Hz). The coherence between each EEG channel and the speech envelope was calculated using MATLAB’s *mscohere()* function. Channel-wise coherence values were then averaged to obtain a participant-specific coherence spectrum. Individual spectra were subsequently averaged within each group (control and dyslexic) to derive group-level coherence profiles. Group differences in coherence across frequencies were evaluated using Bayesian t-tests conducted at each frequency point. This analysis yielded Bayes Factors (*BF_10_*), where *BF₁₀* quantifies the evidence in favour of the alternative hypothesis (*H_1_*) relative to the null hypothesis (*H_0_*). Values greater than 3 indicate substantial evidence for a difference and values between 1 and 3 providing anecdotal or weak evidence favouring the alternative hypothesis. Frequencies yielding *BF_10_* > 3 were considered to show substantial evidence for reduced cerebro-acoustic coherence in one group relative to the other.

### 2.9. Computation of band-power

Neural response power was quantified for each participant and frequency band (delta, theta, beta, and low gamma). Power spectral density (PSD) was first computed for each EEG channel using the *pspectrum()* function in MATLAB, providing a channel-wise PSD as a function of frequency. Channel-level PSDs were then averaged to obtain a broad-band PSD for each trial. For each participant, trial-wise PSDs were further averaged to yield a participant-level PSD representing the overall spectral profile across the entire recording period. Total power within each frequency band was subsequently derived from the participant-level PSDs. Group-level PSDs were computed by averaging individual PSDs across participants within each group (control and dyslexic). This analysis provided estimates of mean neural power for each frequency band and group, enabling comparison of spectral power characteristics across delta, theta, beta, and low-gamma bands.

### 2.10. Computation of power spectral topographic map

Power spectral topographic maps were generated for each frequency band of interest (delta, theta, beta, and low gamma) across the scalp. Outer EEG channels (E17, E48, E119, E125, E126, E127, and E128) were excluded from the topographic maps to minimise edge effects and spatial interpolation artefacts. The PSD was computed over a broad frequency range (0.2–42 Hz) using Welch’s method for each participant and grouped according to participant classification (control or dyslexic). Mean spectral power within each frequency band was then calculated for both groups. Group-level statistical comparisons were performed to identify differences in power between dyslexic and control participants, with particular focus on the right temporal region (EEG channels: E100, E101, E102, E107, E108, E109, E113, E114, E115, E116, E120, E121).

### 2.11. Computation of cross-frequency PAC

Cross-frequency PAC was quantified using the modulation index (*MI*; Tort et al., 2008; Hülsemann, 2019), defined as:

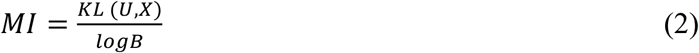

where *B* (=18) denotes the number of phase bins, 𝑈 is the uniform distribution, 𝑋 is the empirical distribution of amplitudes, and 𝐾𝐿 (𝑈, 𝑋) represents the Kullback–Leibler divergence between these distributions. The Kullback–Leibler divergence was computed as:

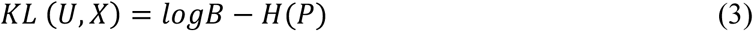

where 𝐻(.) denotes the Shannon entropy, and *P* denotes the vector of normalised averaged amplitudes per phase bin, which is calculated as:

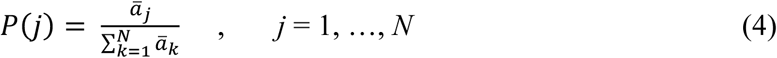

where 𝑎A: is the mean amplitude within the *j*th bin, and *k* serves as the index over all *N* bins. Cross-frequency PAC values were computed for delta-theta, delta-beta, theta-beta, delta-low gamma, and theta-low gamma frequency pairs.

### 2.12. Computation of cross-frequency PPC

Cross-frequency PPC quantifies the degree of phase synchrony between neural oscillations in two different frequency bands. PPC was estimated using the phase-locking value (PLV;

Lachaux et al., 1999), defined as:

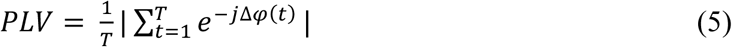

where | . | denotes the absolute value operator, 𝑇 is the number of time samples, and Δ𝜑(𝑡) is the instantaneous phase difference between the two oscillations, calculated as:

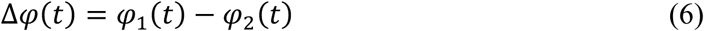

where 𝜑_’_(𝑡) and 𝜑_>_(𝑡) represent the instantaneous phase of first and second signals, respectively. PPC values were computed for five frequency pairs: delta-theta, delta-beta, theta-beta, delta-low gamma, and theta-low gamma.

## 3. Results

### 3.1. Assessing normative speech decoding for dyslexics using between-group analyses

The between-group backward TRF analysis simulated normative decoding by constructing a group-level model from EEG data of control participants, which was then used to test differences in the pattern and accuracy of neural representations between the two groups. Both groups exhibited decoding accuracies significantly above chance for the delta and theta bands, indicating meaningful neural representations of low-frequency envelope information for both groups. Figure S1 illustrates these results, showing that observed decoding performance exceeded chance for both delta (Figure S1A) and theta (Figure S1B) bands.

Figure 3 shows bar plots of envelope reconstruction accuracy for the control and dyslexic groups for the delta (Panel 3A) and theta (Panel 3B) bands, derived from the between-group backward TRF analysis. Independent two-sample t-tests were conducted to assess group differences in decoding accuracy for each frequency band. Participants with dyslexia exhibited significantly lower decoding accuracy than control participants for both the delta (*p* < 0.001) and theta (*p* < 0.001) bands. This is suggestive of impaired neural tracking of both prosodic-rate and syllabic-rate speech information for adults with dyslexia.

**Figure 3.**
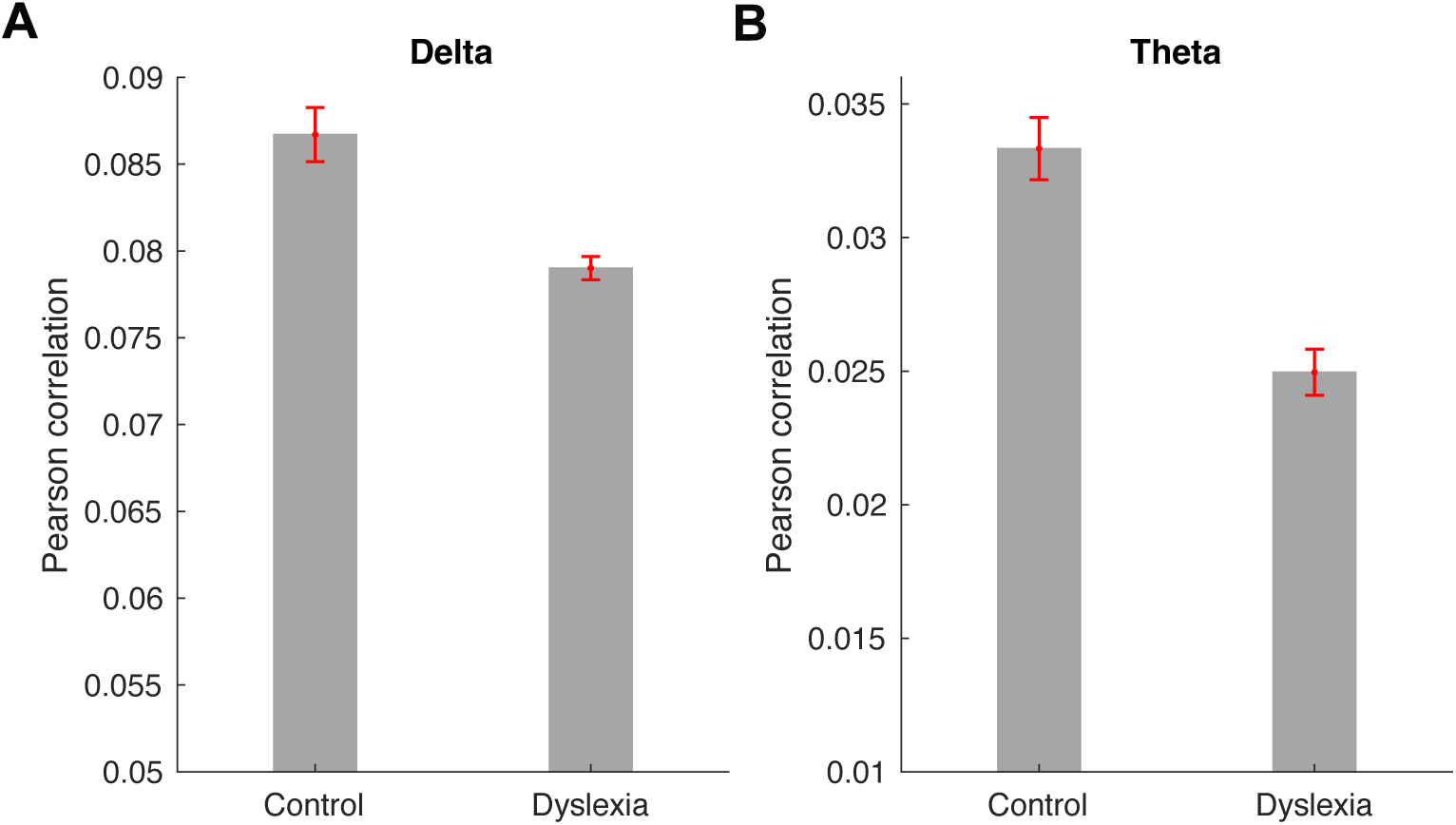
Stimulus reconstruction accuracy for control and dyslexic participants in the delta (Panel A) and theta (Panel B) bands, derived from the between-group analysis. Error bars indicate the standard error of the mean. Note that the y-axis scales differ between panels.

### 3.2. Comparing decoding accuracy across groups using within-participant analysis

The between-group differences described in section 3.1 do not necessarily mean that an individual dyslexic brain has encoded less precise or ‘noisy’ information. It is equally possible that the dyslexic brain is encoding highly precise yet different speech-based representations in comparison to typically developing brains. To test this possibility, the within-participant approach quantified decoding accuracy at the individual level, allowing an estimation of the fidelity of neural speech representations for any given brain. Both groups exhibited significantly above-chance decoding accuracy (*α* = 0.05) for the delta and theta bands, indicating that neural activity reliably represented the low-frequency components of the speech envelope for both groups. Figure S2 shows the PDFs of the null models and the corresponding group-level mean decoding accuracies for the delta (Figure S2A) and theta (Figure S2B) bands. Figure 4 shows box plots of stimulus reconstruction accuracy for the control and dyslexic groups for the delta (Figure 4A) and theta (Figure 4B) bands, derived from the backward TRF models constructed for each participant. These plots illustrate the distribution and variability of decoding accuracy across individuals within each group and frequency band. Independent two-sample t-tests showed a significant group difference for the theta band (*p* = 0.03), adults with dyslexia showing lower decoding accuracy than controls. No significant difference was found for the delta band (*p* = 0.46). This selective deficit in theta-band decoding suggests that adults with dyslexia exhibit diminished neural tracking of syllabic-rate speech information. The result contrasts with findings for children with dyslexia using the same paradigm (Keshavarzi et al., 2022a), where no theta-band difference was observed. The difference for adults may reflect the cumulative impact of reduced reading experience on neural encoding of speech rhythms by the time adulthood is reached.

**Figure 4.**
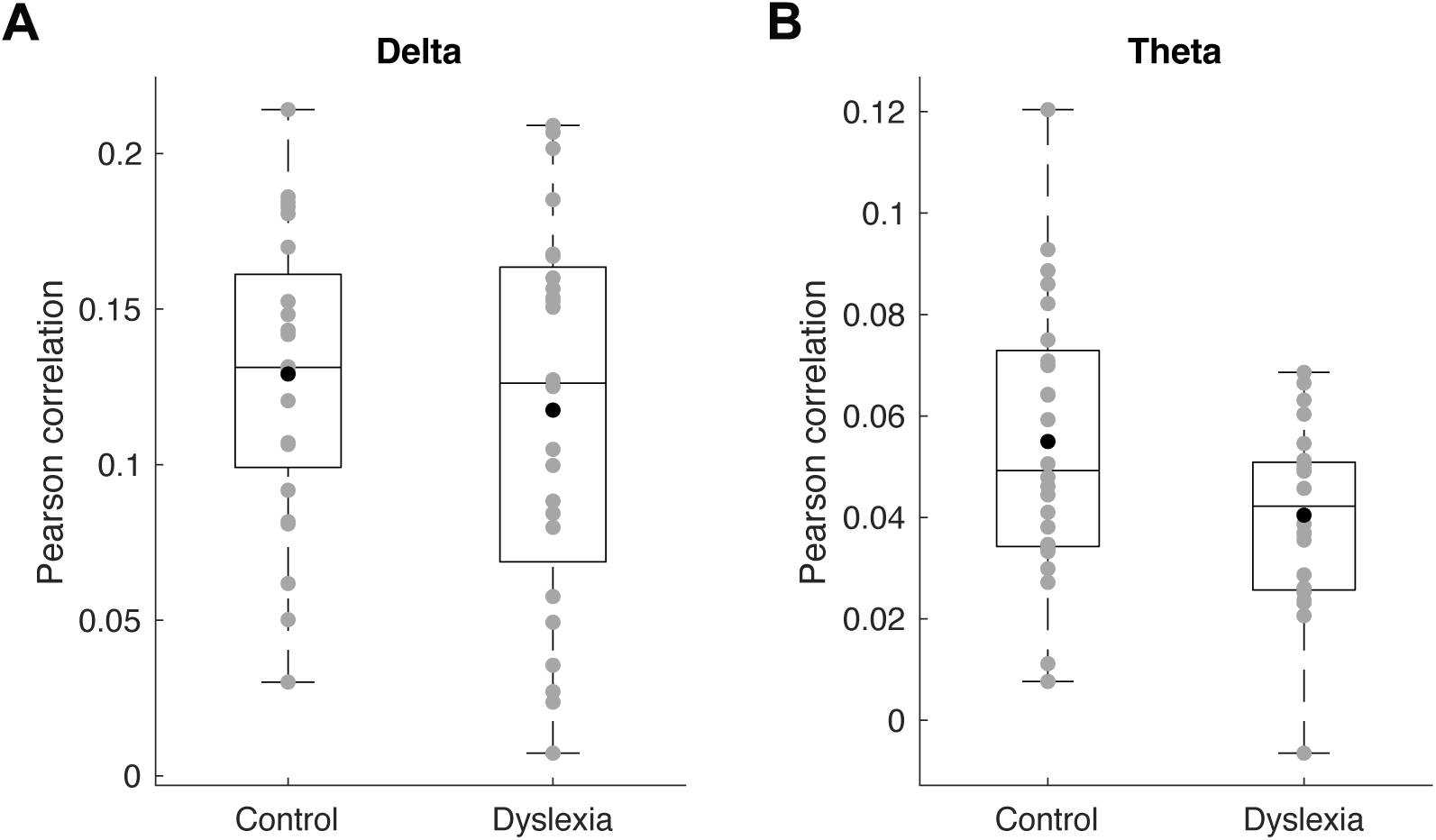
Within-participant stimulus reconstruction accuracy for control and dyslexic participants for the delta (Panel A) and theta (Panel B) bands. Grey circles represent individual reconstruction accuracy scores, and black circles denote group means. The central line in each box indicates the median, the box edges correspond to the 25th and 75th percentiles, and the whiskers extend to the most extreme data points.

### 3.3. Comparing cerebro-acoustic coherence between groups

To examine group-level differences in cerebro-acoustic coherence, we analysed the synchrony between neural activity and the speech envelope separately for the control and dyslexic groups. Coherence was lower for the dyslexic group than for the controls (*p* < 0.05, *BF10* > 3). This was the case for several frequencies within the delta and theta ranges, indicating atypical neural synchronisation with the low-frequency components of continuous speech. Figure 5 presents the coherence spectra for both groups, the control group (blue) showing higher coherence values than the dyslexic group (red). Frequencies with significant group differences are marked by thicker green lines. These findings suggest that dyslexia is associated with impaired low-frequency neural tracking of the temporal dynamics of speech in the frequency range 2–6 Hz, consistent with TS theory. In the current analysis, group differences were largely absent at rates above 6 Hz.

**Figure 5.**
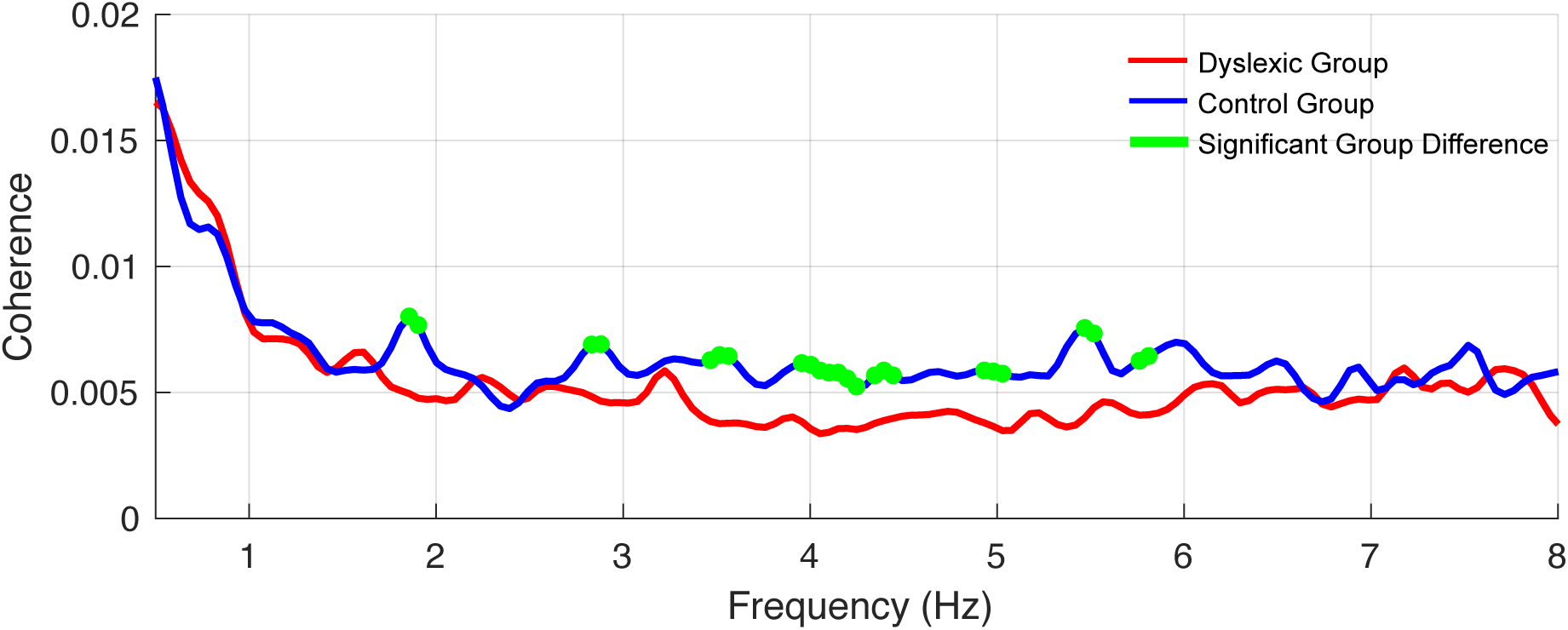
Cerebro-acoustic coherence spectra for control (blue) and dyslexic (red) groups for low frequencies. Thicker green lines indicate frequency regions showing significant group differences (*p* < 0.05, *BF*_10_ > 3).

### 3.4. Comparing band-power between groups across the whole brain

To examine group-level differences in whole-brain neural oscillatory activity, band power values were compared between control and dyslexic participants for four frequency bands: delta, theta, beta, and low gamma. Statistical comparisons were conducted using two-tailed Wilcoxon rank-sum tests. Outliers were excluded prior to analysis; a data point was considered an outlier if its power value exceeded 1.5 times the interquartile range above the upper quartile or below the lower quartile of the group distribution. Mean power values were computed separately for each frequency band and group. A significant group difference was found for the delta band, with greater power for the dyslexic group than for controls (*z* = −2.08, *p* = 0.038; three outliers in the control group, two in the dyslexic group). No significant differences were observed for the theta (*z* = −0.94, *p* = 0.35), beta (*z* = −0.32, *p* = 0.75), or low-gamma (*z* = −0.14, *p* = 0.89) bands. Figure 6 displays band power for each group across the delta (Figure 6A), theta (Figure 6B), beta (Figure 6C), and low-gamma (Figure 6D) frequency bands.

**Figure 6.**
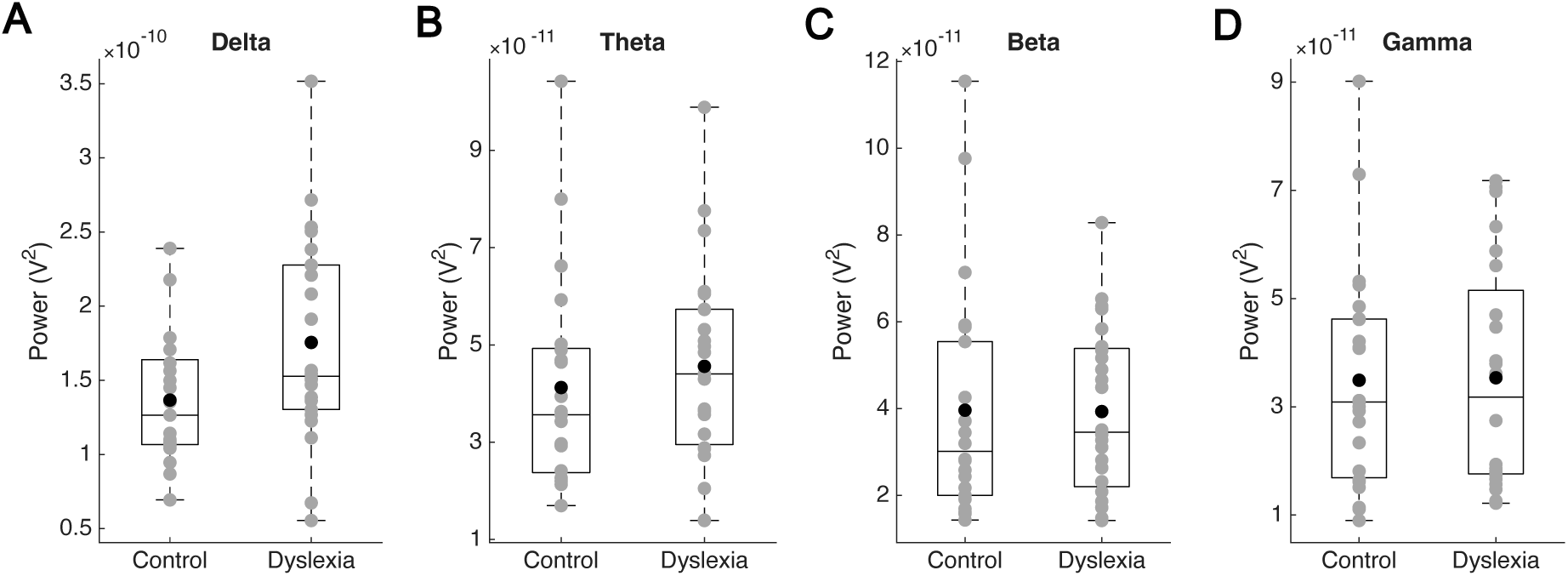
Box plots of power values for the delta (Panel A), theta (Panel B), beta (Panel C), and low-gamma (Panel D) frequency bands. Grey circles represent individual participant values, black circles indicate group means, and horizontal lines within boxes denote medians. Outliers were excluded prior to plotting. Note that the y-axis scales differ across panels.

### 3.5. Comparing spectral power topographic maps between groups

To examine group-level differences in spectral power, topographic maps were generated for each frequency band (delta, theta, beta, and low gamma) and for each group (control and dyslexic). These maps illustrate the spatial distribution of power across the scalp, providing insight into regional variations in neural activity. Figure 7 presents the topographic maps for both groups and the corresponding difference maps, highlighting regions where one group exhibited greater power than the other. Based on prior findings (Arns et al., 2007; Cutini et al., 2016), we hypothesised that delta-band power would be greater in the right temporal region for the dyslexic group than for the controls. This hypothesis was tested using separate two-tailed Wilcoxon rank-sum tests for each frequency band. There was a significant group difference for the delta band (*p* = 0.028), with higher power for the dyslexic group within the right temporal region. No significant differences were found for the theta (*p* = 0.35), beta (*p* = 0.30), or low gamma (*p* = 0.10) bands.

**Figure 7.**
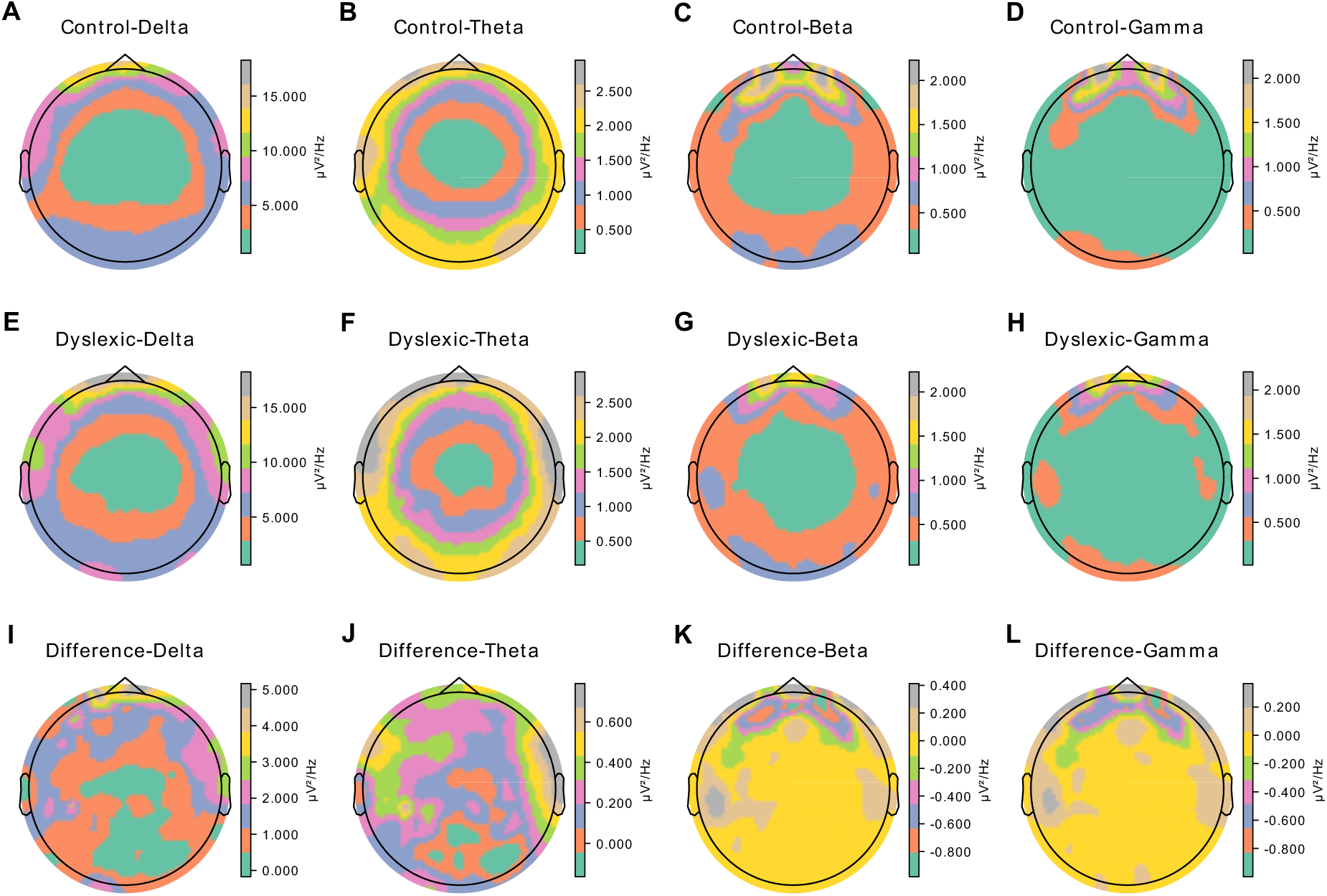
Spectral power topographic maps for control and dyslexic groups and their differences. Panels A–D show delta-, theta-, beta-, and low gamma-band topographic maps, respectively, for control participants. Panels E–H show the corresponding maps for dyslexic participants. Panels I–L depict the subtraction maps (dyslexic minus control), illustrating regions with higher power in one group relative to the other. Colour bars indicate the range of spectral power. Note that the colour scales differ across panels.

### 3.6. Comparing cross-frequency PAC between groups

PAC, particularly theta-gamma PAC, has been shown to play an important role in neurotypical adult speech processing (Giraud & Poeppel, 2012). To examine potential group-level differences in PAC, MI values were computed for five frequency pairs: delta-theta, delta-beta, delta-low gamma, theta-beta, and theta-low gamma. Figure S3 displays the PAC results for both groups across all frequency combinations. No significant group differences were observed for any of the PAC measures: delta-theta (*z* = −0.89, *p* = 0.37; 3 outliers in the control group, 1 outlier in the dyslexic group), delta-beta (*z* = 1.53, *p* = 0.13; 2 outliers in the control group, 1 outlier in the dyslexic group), theta-beta (*z* = 1.32, *p* = 0.19; 1 outlier in the control group, 1 outlier in the dyslexic group), delta-low gamma (*z* = 1.40, *p* = 0.16; 4 outliers in the control group, 2 outliers in the dyslexic group), and theta-low gamma (*z* = 0.97, *p* = 0.33; 4 outliers in the control group, 2 outliers in the dyslexic group). These findings indicate that adults with dyslexia and typical readers exhibit comparable levels of cross-frequency PAC during natural speech processing. Thus, group-level differences in neural oscillatory dynamics, if present, are unlikely to reflect alterations in PAC mechanisms.

#### 3.2.7. Comparing cross-frequency PPC between groups

To examine potential group-level differences in PPC, PLV values were computed for five frequency pairs: delta-theta, delta-beta, delta-low gamma, theta-beta, and theta-low gamma. Figure S4 presents the PPC results for both groups across all frequency combinations. No significant group differences were found for any PPC measure: delta-theta (*z* = −0.20, *p* = 0.84; 1 outlier in the control group), delta-beta (*z* = −0.79, *p* = 0.43), theta-beta (*z* = −0.54, *p* = 0.59; 1 outlier in the control group), delta-low gamma (*z* = 0.38, *p* = 0.70), or theta-low gamma (*z* = 0.81, *p* = 0.42). These results suggest that group-level differences in neural oscillatory dynamics during natural speech processing do not extend to PPC.

## 4. Discussion

To our knowledge, this study is the first to investigate the accuracy of low-frequency cortical tracking of continuous speech by adults with dyslexia. By applying the same decoding method as used with children with dyslexia, we demonstrated that both control and dyslexic adults exhibited above-chance speech envelope decoding accuracy in the delta and theta bands, indicating that neural systems in both groups can meaningfully represent low-frequency speech envelope information. This contrasts with previous findings for children with dyslexia using the same continuous speech paradigm (Keshavarzi et al., 2022a), where theta-band decoding for dyslexic children in the between-group analysis fell below chance level. The between-group analysis here revealed differences in the pattern of neural representation for both the delta and theta bands, indicating that adults with dyslexia process low-frequency information in continuous speech differently from typical readers. The delta-band data match our findings for dyslexic children (Keshavarzi et al., 2022a), which suggests that atypical delta-band processing of continuous speech persists into adulthood. In the within-participant analysis, delta-band decoding accuracy was comparable for the two groups, indicating that delta-band entrainment during natural speech processing remains intact in adults with dyslexia at the individual level. However, theta-band decoding accuracy was significantly lower for the dyslexic group than for the controls in the within-participant analyses. This differs from previous findings in children (Keshavarzi et al., 2022a), where no group differences for the theta band were observed in the within-participant analyses. This may indicate that cumulative differences in reading experience affect the accuracy of theta-band encoding by adulthood. Adults with dyslexia, who typically engage less frequently in text-based reading, may experience reduced neural tuning to syllabic-rate speech rhythms, leading to weaker theta-band entrainment. These findings suggest that theta-band decoding may be particularly sensitive to experience-dependent plasticity, potentially reflecting prolonged interaction between speech processing and reading development across the lifespan. It is worth noting that adult cortical tracking studies of individuals without dyslexia typically test university students (Giraud & Poeppel, 2012), thereby utilising highly literate populations. The dominance of theta-band cortical tracking in this literature may therefore be confounded with the high reading efficiency of the student populations being tested.

Cerebro-acoustic coherence analyses revealed significantly lower coherence for the dyslexic group than for the controls in the range 2–6 Hz. This is consistent with the findings for Spanish dyslexic participants of Molinaro et al. (2016), who reported reduced coherence at ∼2 Hz. Lower coherence likely indicates diminished synchrony between neural activity and the low-frequency components of the speech envelope, reflecting impaired temporal tracking of prosodic- and syllabic-rate speech information by those with dyslexia. As Molinaro et al. (2016) observed reduced delta-band synchrony for both dyslexic adults and dyslexic children, these findings are consistent with the view that atypical neural synchrony to the prosodic and rhythmic features of speech persists into adulthood. These results are also consistent with TS theory, which posits that dyslexia is characterised by atypical neural entrainment to the low-frequency temporal structure of speech from infancy onwards. The presence of coherence deficits across both delta and theta bands highlights their potential as neurophysiological markers of dyslexia and possible targets for remediation.

A critical distinction between the present findings and those of Molinaro et al. (2016) concerns both the nature of the speech input and the level at which cortical tracking was quantified. Molinaro et al. demonstrated reduced delta-band cerebro-acoustic coherence in dyslexic adults during sentence listening, with no evidence for atypical coherence in the theta band. Using a substantially longer continuous narrative and combining coherence with reconstruction-based decoding, the present study extends these findings in two important ways. First, we show that atypical low-frequency (delta and theta) tracking persists during sustained, ecologically valid speech comprehension, not only during brief sentence-level processing. Notably, group differences were observed in both frequency bands when assessed using cerebro-acoustic coherence, indicating broader low-frequency synchronisation deficits than previously reported in adults. Second, reduced decoding accuracy – particularly in the theta band – demonstrates that dyslexic adults do not merely exhibit weaker synchronisation, but also encode syllabic-rate speech information with reduced representational precision. This convergence across coherence- and decoding-based metrics strengthens the interpretation that atypical theta-band processing reflects altered neural representation rather than a measurement-specific effect. This distinction is important, as reduced coherence alone cannot determine whether neural representations are noisy, systematically altered, or simply phase-misaligned. Our findings indicate that atypical low-frequency speech encoding in adult dyslexia reflects both altered synchronisation and diminished representational fidelity.

The band-power analysis revealed significant group differences in the delta band across the whole brain, with higher power for the dyslexic group than for controls. Against our expectations, no significant group differences were found for the beta band. Topographic analyses corroborated the delta-band findings, showing increased power in the right temporal region of dyslexic participants relative to controls. These right-lateralised findings are consistent with AST theory, suggesting atypical right-lateralised oscillatory neural processing in dyslexia over longer temporal integration windows (150–250 ms, Poeppel, 2003). The present delta-band power findings are consistent with data for the rhythmic audiovisual speech paradigm reported by Keshavarzi et al., (2025), which was applied to the same adult participants as tested here. In the study of Keshavarzi et al., the dyslexic participants also exhibited significantly increased delta-band power both across the whole brain and specifically within the right temporal region. This spatial pattern is consistent with TS theory and with previous reports by Arns et al. (2007) and Cutini et al. (2016), which similarly identified atypical right-hemispheric activity in children with dyslexia. These findings suggest that enhanced delta-band power in the right temporal region may serve as a neural marker of dyslexia, potentially reflecting compensatory or inefficient processing mechanisms during speech perception.

Neither PAC nor PPC analyses revealed significant differences between the control and dyslexic groups across any of the frequency pairs examined (delta-theta, delta-beta, delta-low gamma, theta-beta, and theta-low gamma). These results indicate that cross-frequency interactions, as indexed by PAC and PPC, do not appear to account for the group-level differences in neural dynamics observed during natural speech processing. It is possible that such coupling mechanisms may emerge under different task demands or reflect other aspects of speech and language processing not captured by the present paradigm. Further research using complementary paradigms and multimodal approaches will be valuable for clarifying the role of cross-frequency coupling in dyslexia.

The findings of this study are broadly consistent with the TS theory, a developmental framework that attributes phonological deficits in dyslexia to impaired neural entrainment in low-frequency oscillatory bands from infancy onwards, particularly delta and theta. Several of the current findings, including reduced theta-band decoding accuracy in the within-participant analysis, group-level differences in the neural representation patterns for both delta and theta bands, and decreased cerebro-acoustic coherence in these same frequency ranges, collectively support TS theory’s emphasis on the role of low-frequency oscillations in developmental dyslexia. The data presented here suggest that atypical low-frequency decoding does not disappear with age or with reading experience. The reduction in theta-band decoding accuracy, which was observed for dyslexic adults but not children, may indicate that reading experience influences oscillatory encoding over developmental time. Rhythmic training or neurostimulation approaches that enhance neural synchronisation to speech rhythms at low-frequency temporal rates (Riecke et al., 2018; Keshavarzi et al., 2020) administered early in development could thus help to improve phonological and reading outcomes for individuals with dyslexia. Improving low-frequency neural synchrony using biofeedback or other approaches may also represent a promising pathway for remediation.

## Author Contributions

M.K. conceptualisation, methodology, data collection, data analyses, visualisation, writing-original draft; B.C.J.M. conceptualisation, methodology, writing-review & editing; U.G. funding acquisition, project administration, supervision, conceptualisation, methodology, writing-original draft.

## Data Sharing and Data Availability

De-identified data and analysis code are available from the corresponding author upon reasonable request, subject to institutional and ethical constraints.

## Acknowledgements

The authors would like to thank all the participants involved in the study. This research was funded by a donation to UG from the Yidan Prize Foundation. The sponsor played no role in the study design, data interpretation nor writing of the report.

## Supplementary Information

**Figure S1.**
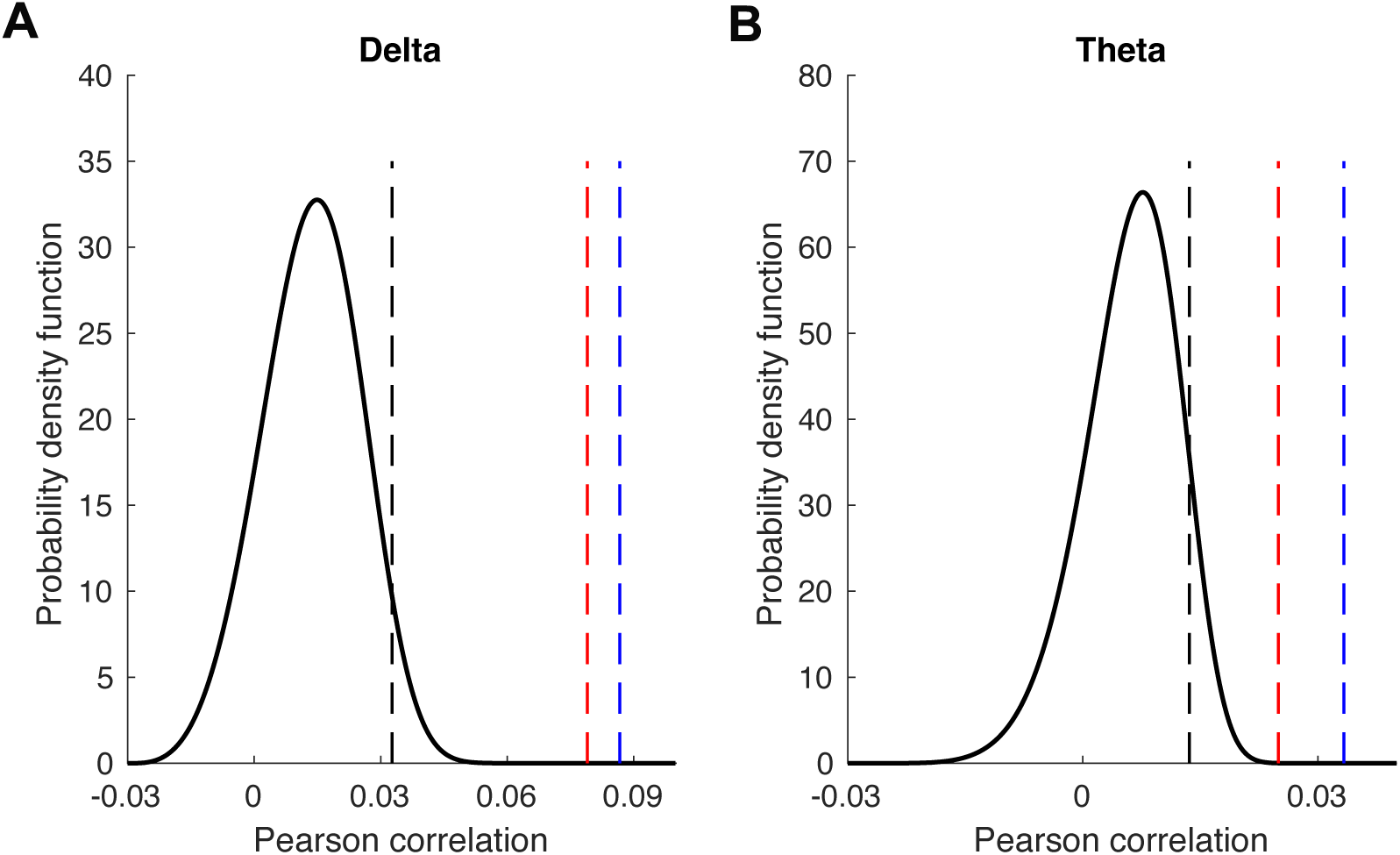
Statistical evaluation of reconstruction accuracy for the between-group analysis of the delta (Panel A) and theta (Panel B) bands. Blue and red lines represent the control and dyslexic groups, respectively. Black curves show the probability density functions (PDFs) of the null models, defining chance-level performance. Dashed black lines indicate correlation thresholds required for statistical significance (*α* = 0.05), while dashed coloured lines denote the mean correlation (a measure of reconstruction accuracy) for each group and each band. Note that the x-axis scales differ between panels.

**Figure S2.**
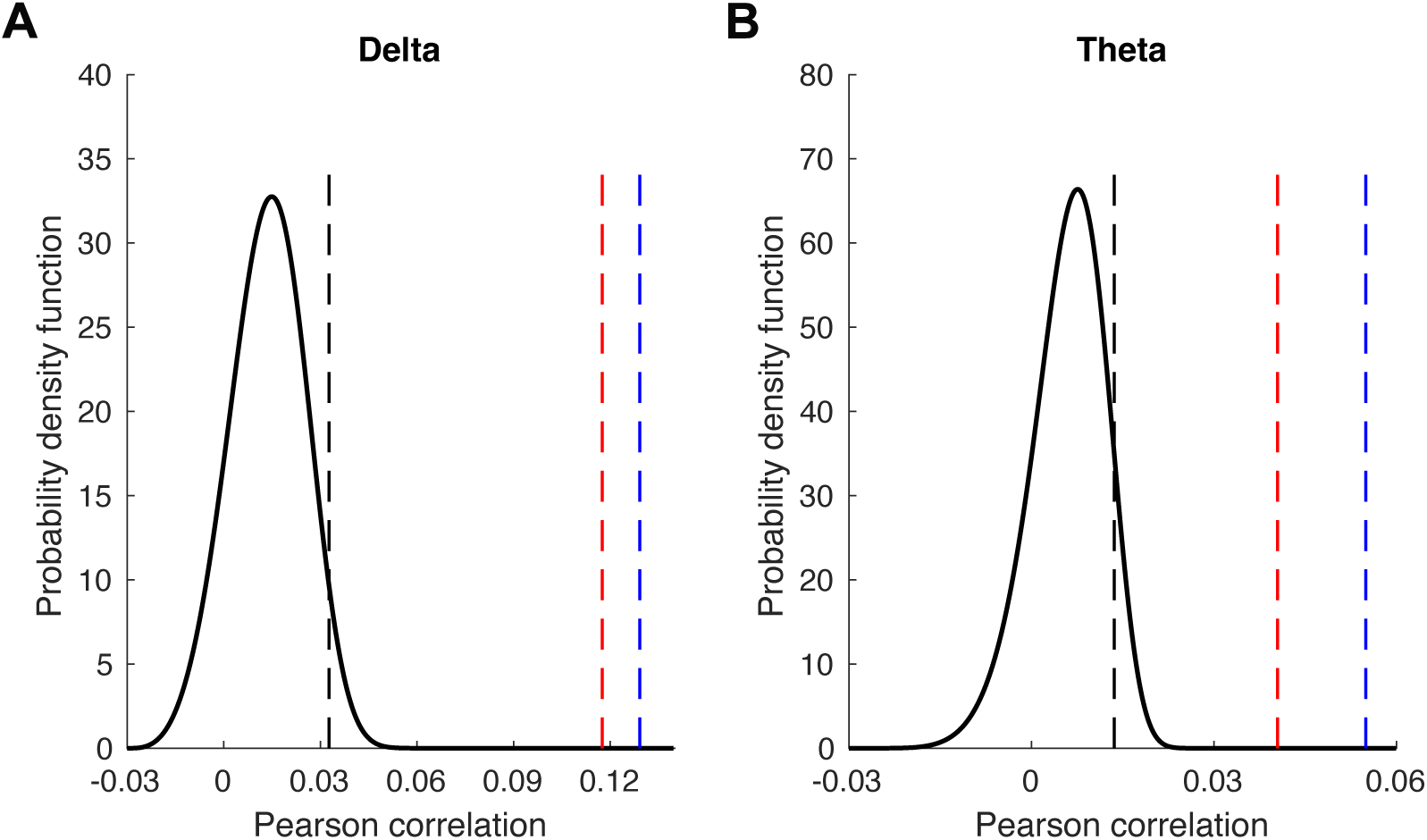
Statistical assessment of reconstruction accuracy for the within-participant analysis of the delta (Panel A) and theta (Panel B) bands. Blue and red lines represent the control and dyslexic groups, respectively. The black curves show the probability density functions (PDFs) of the null models, defining chance-level performance. Dashed black lines indicate correlation thresholds required for statistical significance (*α* = 0.05), while dashed coloured lines denote the mean correlation (a measure of reconstruction accuracy) for each group and each band. Note that the x-axis scales differ between panels.

**Figure S3.**
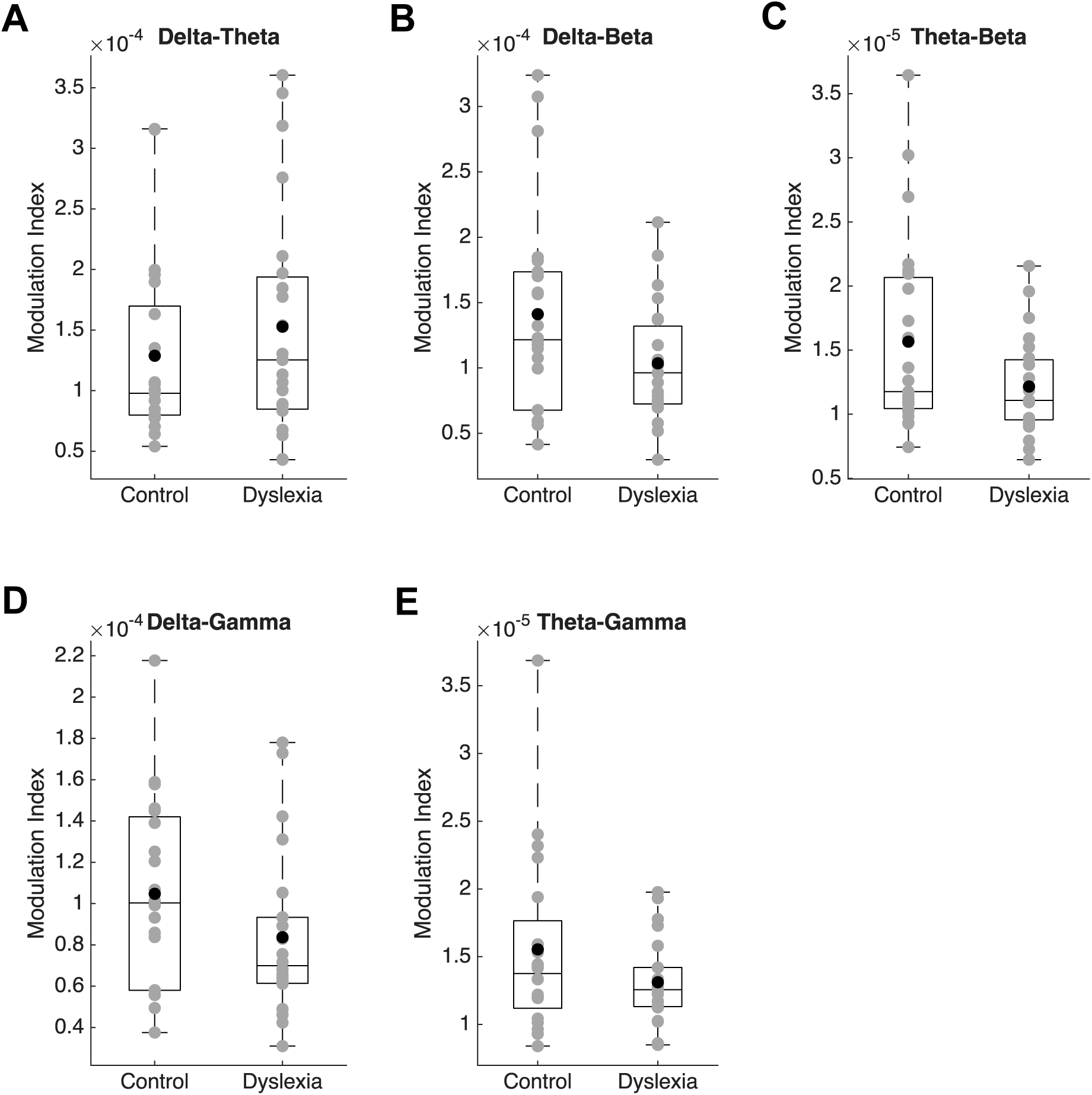
Box plots of PAC values (Modulation index) for the delta-theta (Panel A), delta-beta (Panel B), theta-beta (Panel C), delta-low gamma (Panel D), and theta-low gamma (Panel E) frequency pairs. Grey circles represent individual participant values, black circles indicate group means, and horizontal lines within boxes denote medians. Outliers have been excluded.

**Figure S4.**
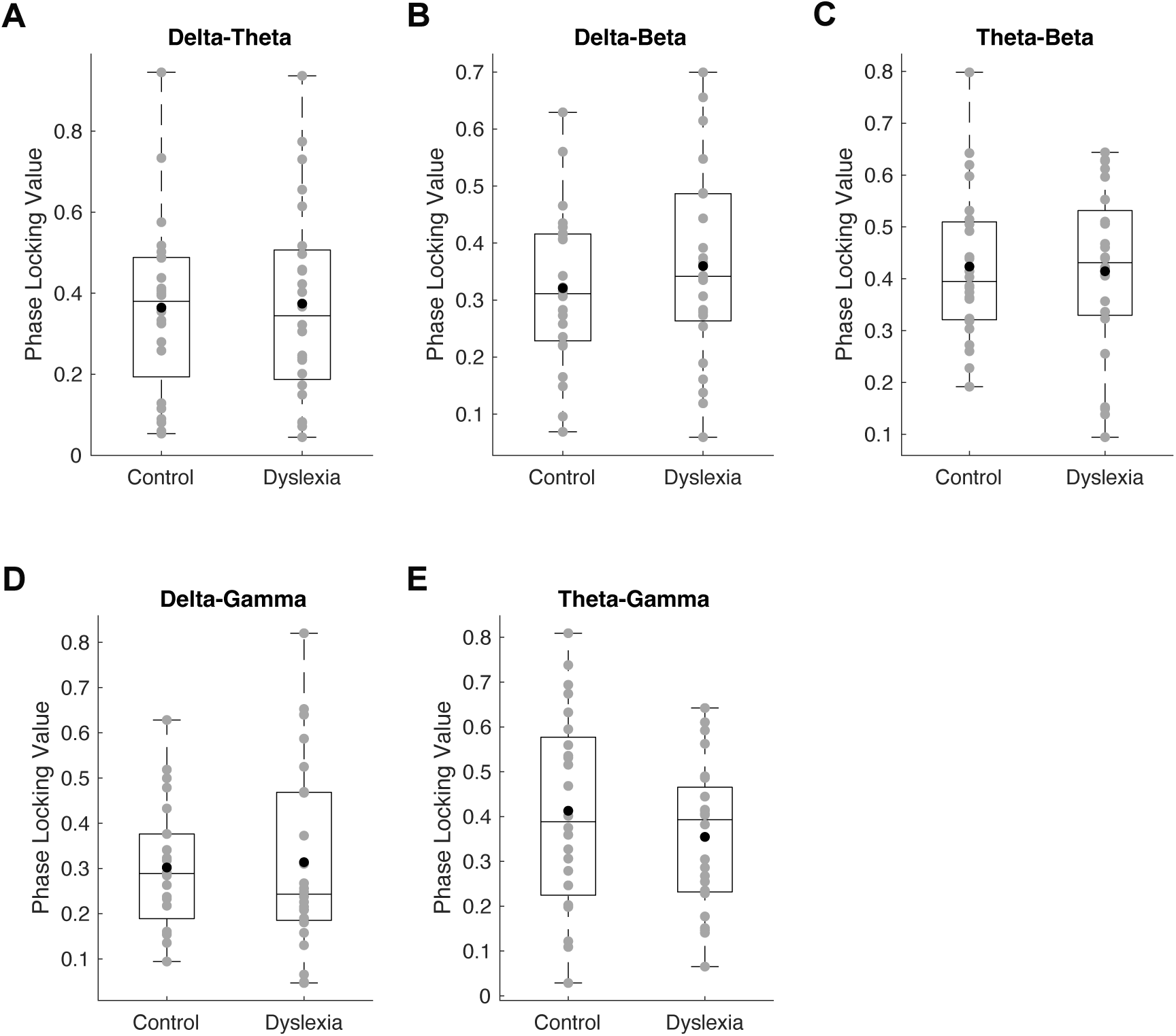
Box plots of PPC values (phase locking values) for the delta-theta (Panel A), delta-beta (Panel B), theta-beta (Panel C), delta-low gamma (Panel D), and theta-low gamma (Panel E) frequency pairs. Grey circles represent individual participant values, black circles indicate group means, and horizontal lines within boxes denote medians. Outliers have been excluded.

